# Interpreting turbidity measurements for vesicle studies

**DOI:** 10.1101/348904

**Authors:** Anna Wang, Christopher Chan Miller, Jack W. Szostak

**Affiliations:** Department of Molecular Biology and Center for Computational and Integrative Biology, Massachusetts General Hospital, Boston MA 02114, USA; Atomic and Molecular Physics Division, Harvard Smithisonian Center for Astrophysics, Cambridge MA 02138, USA

**Author notes:** Electronic mail (Anna Wang). Electronic mail (Jack W. Szostak).

## Abstract

Spectrophotometers are routinely used to assess the turbidity of vesicle solutions. Here we provide guidelines for interpreting turbidity measurements of vesicle samples, and highlight potential pitfalls of the approach. We use an exact solution for core-shell scatterers to model and calculate how samples of vesicles scatter light, and provide a comprehensive overview of how the turbidity of vesicle samples can change with vesicle size, contents, and composition. Surprisingly, we find that vesicle lamellarity has a large effect on sample turbidity, while unilamellar vesicles of different sizes have similar turbidity. We use our model in conjunction with experimental data to measure the thickness of oleic acid vesicle membranes and find excellent agreement with values determined by cryo-TEM. We also calculate the effects of potential errors in measurement from forward scattering and multiple scattering.

## I. INTRODUCTION

The desire to characterise and monitor the properties of vesicles - semi-permeable membranes that enclose an aqueous compartment (Fig. 1a) - comes from many fields. Vesicles can be biocompatibile and deliver cargo for drug deliver^1,2^, are used as compartments for synthetic biology^3^ and origins of life studies^4^, and are used as model systems to study cell-membrane properties^5^. A non-invasive way to determine the properties of vesicles is to use their interaction with visible light. For example, dynamic light scattering (DLS) is often used for vesicle sizing^6^, microscopy is used to determine morphologies of vesicles^7^, multiangle light scattering is used during flow cytometry to distinguish between cell types^8^, and fluorescence measurements are routinely used to assess vesicle encapsulation efficiency^9^.

**FIG. 1.**
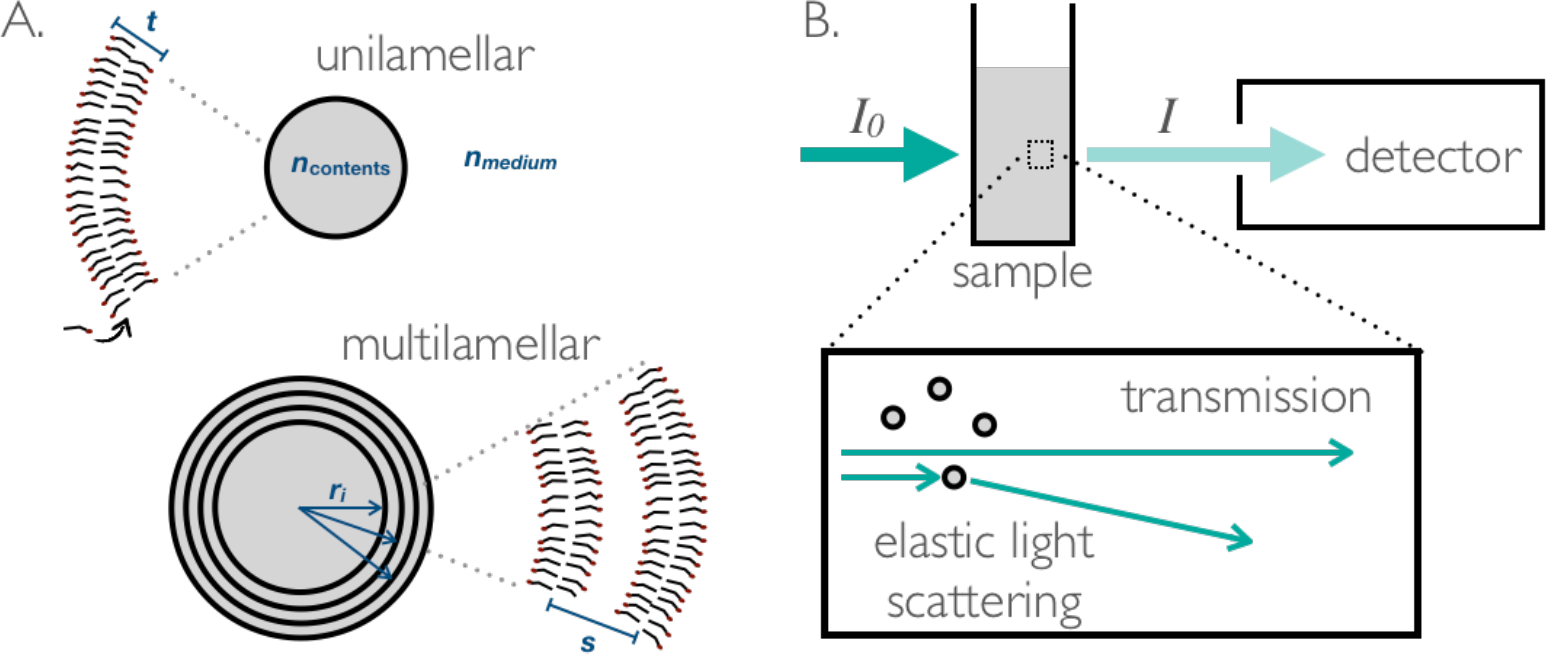
**A**. Vesicles are semi-permeable compartments delineated by a membrane. The membrane is typically a bilayer of amphiphile, and encloses a material of refractive index *n*_contents_ in a medium of refractive index *n*_medium_. Unilamellar vesicles have one membrane of thickness *t*, and multilamellar vesicles have many membranes with centre-to-centre separations *s*. **B**. The turbidity of vesicle samples can be measured on a spectrophotometer. The a detector measures the intensity of light *I* that passes 180° through a sample with an illumination source of intensity *I*_0_. This is typically a measure of how much light is unattenuated (transmitted).

One particularly simple but versatile optical technique is turbidimetry (Fig. 1B), where the turbidity of a sample is measured using the widely-available spectrophotometer. Researchers have used this technique to assess a variety of vesicle properties, including vesicle formatio^10–12^, dissolution^13^, permeability^14^, flocculation^15^, average vesicle siz^16–19^, membrane bilayer thickness^20^, and change in average vesicle size^21^. Changes in turbidity are usually attributed to just a single parameter, even though the turbidity of a sample depends on all of these quantities.

It is precisely the versatility of turbidimetry that begs the following question: if the turbidity of a sample increases or decreases, how is it possible to determine which vesicle property is causing the change? Some papers invoke light scattering theory to help tease apart these effects, and often use the Rayleigh-Gans-Debye approximatio^16,17,20^ to achieve an analytical form. However, the approximation begins to break down when vesicles encapsulate materials that differ from the outside, or when vesicles have multiple bilayers, otherwise known as multilamellar vesicles (Fig. 1A).

Here we use a multilayered-sphere light-scattering solution^22,23^ that explicitly specifies the bilayers and inter-bilayer spaces to calculate how vesicles scatter light. This model can handle encapsulated contents, multilamellarity, and larger scatterers than the Rayleigh-Gans-Debye approximation is suited for. By relating a scattering cross section to sample turbidity, we show how sample turbidity depends on vesicle size, composition, contents, and lamellarity. As a guide for when quantitative turbidity measurements can be meaningfully made, we also show the scattering phase functions - how light scatters as a function of direction - for various vesicle types, and discuss the effects of significant forward and multiple scattering.

We then present two contrasting examples of how turbidity measurements can be used for vesicle studies. We first demonstrate potential pitfalls when interpreting experimental turbidity measurements. We then measure the bilayer thickness of oleic acid vesicles to excellent agreement with literature, showing that when used in conjunction with dynamic light scattering, optical microscopy, and light scattering calculations, turbidity can be a quantitative and powerful tool.

## II. MODEL

### A. Measuring turbidity on a spectrophotometer

We first seek to relate how individual vesicles scatter light to the measured turbidity of a vesicle sample. In general, there are three main outcomes for a photon as it encounters a vesicle (Fig. 1): it can pass by without interacting (be transmitted), be *absorbed* (for example, by a fluorophore or a dye), or elastically *scatter* off the vesicle without changing its wavelength^24^.

Once a photon passes through an entire sample, there are analogous quantities that describe the effect of the whole sample on light. The *extinction ∊* refers to the attenuation of photons as they pass through a sample, and consists of the attenuation owing to absorption (*absorbance A*) and the attenuation owing to scattering (*turbidity τ*). In this study we mainly consider samples in which a photon will not interact with more than one vesicle as it travels through a sample. This is known as the *single scattering* regime. Experimentally, a sample is in this regime if the extinction scales linearly with the sample concentration. We will also briefly discuss the effects of multiple scattering and scattering in the direction of the detector.

Decades ago, it was proposed that spectrophotometers can be used to measure not just the absorbance of absorbing samples, but also the turbidity of non-absorbing samples^16^. This is because while spectrophotometers report an ‘absorbance’ *A*, the quantity that is actually measured is how much light does *not* make it through the sample towards a detector situated opposite from the light source (see Fig. 1B): the extinction *∊*.

Thus a typical spectrophotomer reports an ‘absorbance’ as follows:

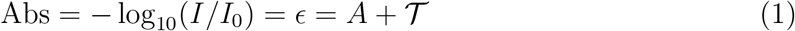

where *I*_0_ is the incident light intensity, *I* is the intensity of light that enters the detector, 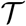 is the extinction owing to scattering. It is assumed that no light is scattered into the detector. We will revisit this assumption in the section examining the effects of scattering towards the detector.

For absorbing samples that have insignificant light scattering 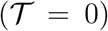, the measured quantity *∊* equals a true absorbance *A* and the concentration of a sample can be determined by using *A* = *∊_A_cl*, otherwise known as the Beer-Lambert law. Here *∊*_*A*_ is the molar absorption coefficient ([*∊*_*A*_] = M^−1^ cm^−1^), *c* is the concentration of the absorbing molecule ([*c*] moles/L = M) and *l* is path length (usually through a cuvette, *l* ~ 1 cm).

When describing non-absorbing samples, the total attenuation of light after it passes through a sample is usually described by a *turbidity* or optical depth τ:

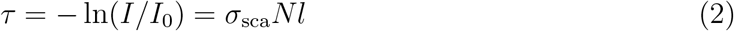

assuming the fraction of light scattered in the direction of the detector is insignificant. *σ*_sca_ is the scattering cross section per scatterer (e.g. a vesicle, [*σ*_sca_] = m^2^), and *N* is the number density of scatterers ([*N*] = m^−3^). By equations 1 and 2, the turbidity of a non-absorbing sample (*A* = 0) can be measured on a spectrophotometer: the ‘absorption’ ∊ measured on spectrophotometers is in fact linearly proportional to the turbidity 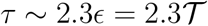.

We now seek to find a molar turbidity coefficient *∊*_τ_ (M^−1^ cm^−1^) analogous to the molar absorption coefficient ∊_A_ in equation 1 to more easily relate calculated and experimental quantities. For a fixed concentration of amphiphile in a system, *c*, the number density of vesicles *N* ([*N*] = m^−3^) is given by

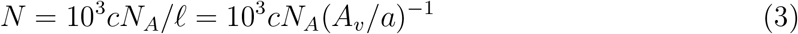

where *N*_*A*_ is Avogadro’s number, ℓ is the number of amphiphile molecules per vesicle, *a* is the area per amphiphile in a membrane, and *A*_*ν*_ is the total amphiphile surface area per vesicle. For example in vesicles where the amphiphiles arrange themselves in bilayers, if a leaflet has *j* amphiphiles then *A*_*ν*_ = 2*ja*.

The turbidity from equation 2 then becomes

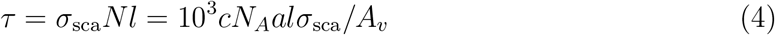

Because the turbidity scales linearly with the concentration of amphiphile *c* and the path length *l*, we can define the molar turbidity coefficient

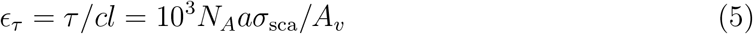

We calculate the scattering cross section *σ*_sca_ with Yang’s recursive algorithm within the light scattering package HoloPy. Equation 4 is then used to calculate the sample turbidity τ.

For absorbing samples, we can modify Equation 4 and calculate

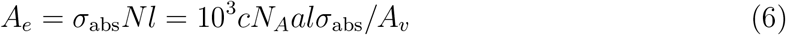

where the absorption cross section *σ*_abs_ appears instead of *σ*_sca_. We can then calculate the absorbance *A* as measured on a spectrophotometer, *A* ~ 2.3*A*_*e*_.

### B. Calculating the scattering cross-section

In general, the scattering cross-section *σ*_sca_ of a micrometer-scale homogeneous particle in a medium with refractive index *n*_medium_ depends on its size, shape and refractive index *n*_particle_. For core-shell structures such as vesicles, the scattering cross-section is much more complex, and depends on the parameters

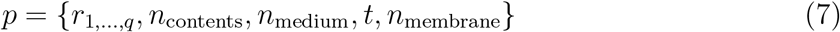

where *r*_*i*_ are the radii of each of the *q* membranes in the vesicle (measured from the centre of the vesicle to the middle of the membrane), *n*_contents_ is the content refractive-index, *n*_medium_ is the external solution refractive-index, *t* is the membrane thickness, and *n*_membrane_ is the membrane refractive-index. In this work we do not consider vesicles with non-concentric centres. We assume a spherical geometry and that the membranes are evenly spaced with *s* being the centre-to-centre spacing between membranes. Though in general the spacing can be arbitrary, for *s* and *t* much smaller than the wavelength of light we do not expect the exact spacing *s* to play a large role in determining turbidity. We define the radius of the vesicle *r* as the distance between the vesicle centre and the centre of the outermost membrane.

For simplicity and ease of comparison to experiments, we report calculated turbidities τ and absorbances A for samples with 5 mM total membrane lipid and a path length of 1 cm unless stated otherwise. The lipid parameters we use are that of a lipid similar to oleic acid, with *n*_membrane_ = 1.46, the bilayer thickness *t* = 3.2 nm, and the area per lipid *a* = 0.311 nm^2^ taken from Han^25^.

## III. RESULTS

### A. Membrane properties: thickness and refractive index

For non-absorbing samples, we first consider how the optical properties of the membrane - its thickness and refractive index – affect the sample turbidity.

To intuitively understand the coupling between membrane thickness and refractive index, we look at one membrane of thickness *t* (Fig. 1) and refractive index *n*_membrane_. Because the thickness *t* is typically a few nanometers, at least two orders of magnitude smaller than optical wavelengths, a membrane and its immediate surroundings of thickness *T* can obey effective medium approximations such as the volume-weighted effective refractive index rule where the effective refractive index of a region is the weighted sum of the volume of each component^26^:

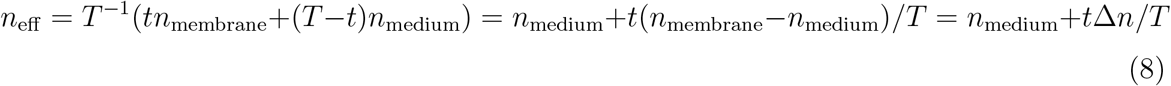

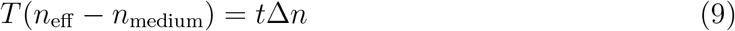

A membrane with index *n*_membrane_ and thickness *t* will therefore interact with light of wavelength λ in the same manner as a membrane with index *n*_*eff*_ and thickness *T*, as long as *t* << λ and *T* << λ. This is because they represent the same optical path difference. A further implication of the bilayer being much thinner than optical wavelengths is that a vesicle with membrane thickness 2*t* will scatter light in the same manner as a vesicle with two closely-spaced membranes (*s* << λ). We will explore scattering from multilamellar vesicles further below.

We find that increasing membrane thickness and refractive index increases sample turbidity (Fig. 2A) and that these curves collapse onto a straight line when plotted against the square of the optical path difference *t*Δ*n* (Fig. 2B). One consequence of this result is that the two parameters *t* and *n* can not independently be determined using turbidity alone.

**FIG. 2.**
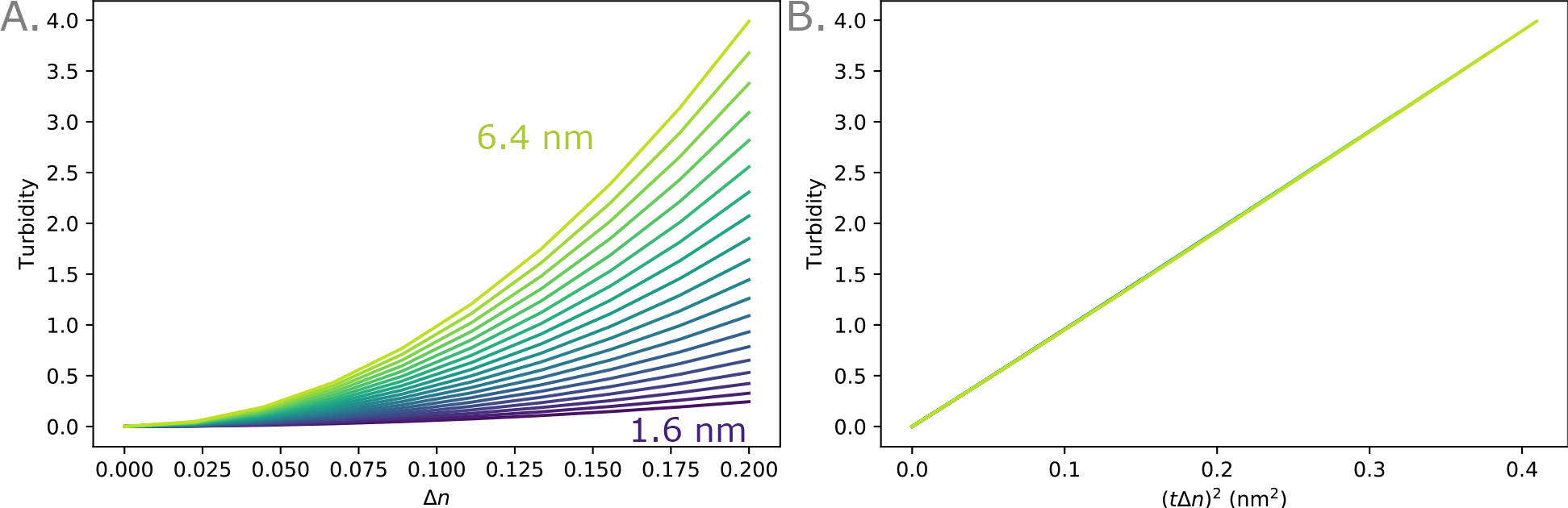
**A**. Calculated turbidity of 5 mM samples of 100-nm-diameter vesicles of increasing membrane thickness *t*. λ = 400 nm. **B**. The calculated turbidity collapses onto a single line when plotted against (*t*Δ*n*)^2^, consistent with refractive index mixing rules (Eq. 9). λ = 400 nm.

### B. Encapsulated content refractive index

Next we consider the effect of any encapsulated vesicle contents on vesicle scattering. Experimentalists routinely use purification methods such as size exclusion chromatograph^27,28^ or dialysis^29^ to remove unwanted solutes that are not encapsulated inside vesicles. Sometimes these solutes are non-absorbing, such as salts, and contribute to the real part of the refractive index Re(*n*_contents_) = *n*; other times they absorb light, and contribute to the imaginary part of the refractive index Im(*n*_contents_) = *k*. Here for simplicity we assume that the medium is non-absorbing, Im(*n*_medium_) = 0, and do calculations at one wavelength (λ = 400 nm). Both *n* and *k* usually vary with wavelength.

For vesicles that encapsulate a non-absorbing solution that has a different refractive index from the surrounding medium (Δ*n*_io_ ≠ 0), the turbidity of the sample increases non-linearly with the vesicle radius (Fig. 3A). Because the surface area to volume ratio of the vesicles decreases with vesicle size, the contribution of even a small Δ*n*_io_ = *n*_contents_ – *n*_medium_ to sample turbidity can easily surpass that of the membrane for larger vesicles. We find that typical values of Δ*n*_io_ are 0.0025 for vesicles with 100 mM encapsulated sucrose and 100 mM glucose in the external aqueous phase, 0.001 for 15 mM encapsulated adenosine 5’- monophosphate (disodium salt), and 0.001-0.002 for 1 mM encapsulated 12-16 long RNA sequences.

**FIG. 3.**
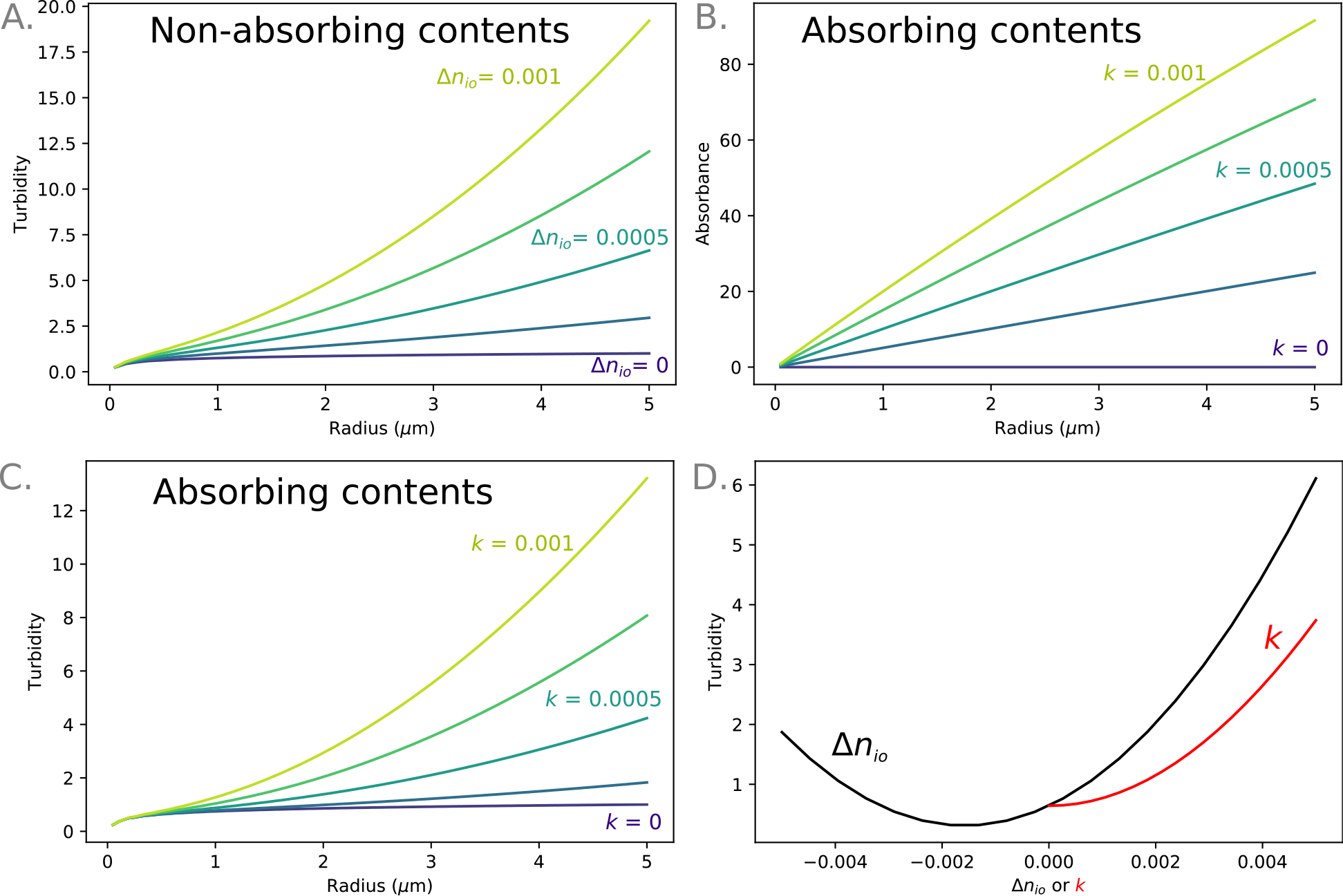
**A**. The turbidity of a non-absorbing sample increases with the refractive index contrast (Δ*n*_io_) between the inside and the outside of the vesicles. λ = 400 nm. **B**. The absorbance *A* of a 5 mM sample of vesicles encapsulating an absorbing material (such as a dye) increases with vesicle radius and *k*. *λ* = 400 nm. **C**. The turbidity of a 5 mM sample of vesicles encapsulating an absorbing material (such as a dye) increases with vesicle radius and *k* (λ = 400 nm). **D**. The turbidity of a 5 mM sample of vesicles increases when either the real or imaginary parts of the refractive index of the contents differ from the refractive index of the medium. λ = 400 nm, *r* = 500 nm.

Using Equation 6, we find that for vesicles encapsulating a solution that absorbs light (*k* ≠ 0), such as a dye, the absorbance of the sample increases linearly with vesicle radius when the total concentration of lipid in the sample is fixed (Fig. 3B). This is because, assuming negligible scattering, the absorbance of the sample increases linearly with the number of absorbing molecules in the sample (Beer’s law) and when the total lipid concentration in a sample is fixed, the encapsulated volume increases linearly with radius (while the number of vesicles decreases).

Interestingly for large enough *k*, encapsulating an absorbing solution also enhances the sample turbidity (Fig. 3C). The measurement for *A* on a spectrophotometer will therefore be affected by ∊_τ_ although the magnitude of the effect is complex and depends on the exact conditions^30^. We expect this effect to be pronounced for vesicles encapsulating a very high concentration of dye, such as self-quenching concentrations of calcein (approximately 100 mM) when monitoring dye leakage. We estimate that *k* is approximately 0.0007 for 1 mM calcein, and 0.02 for 1 mM phycoerythrin by comparing Equation 1 to molar extinction coefficients^31^.

In general, the effect of any refractive index mismatch between the encapsulated contents and medium will contribute to sample turbidity (Fig. 3D). The sample turbidity can even be decreased by an index mixmatch if a lower content index compensates for the higher index membrane.

### C. Vesicle size

Next we consider how the size of vesicles affects sample turbidity. This can be useful for sizing vesicles or monitoring their growth and division. We find that for vesicles where the contents have no refractive index contrast with the medium *n*_contents_ = *n*_medium_, the turbidity of the system increases roughly with the logarithm of vesicle size for a fixed total concentration of lipid in the sample (Fig. 4). This scaling is in contrast to vesicles where *n*_contents_ ≠ *n*_medium_ (Fig. 3).

**FIG. 4.**
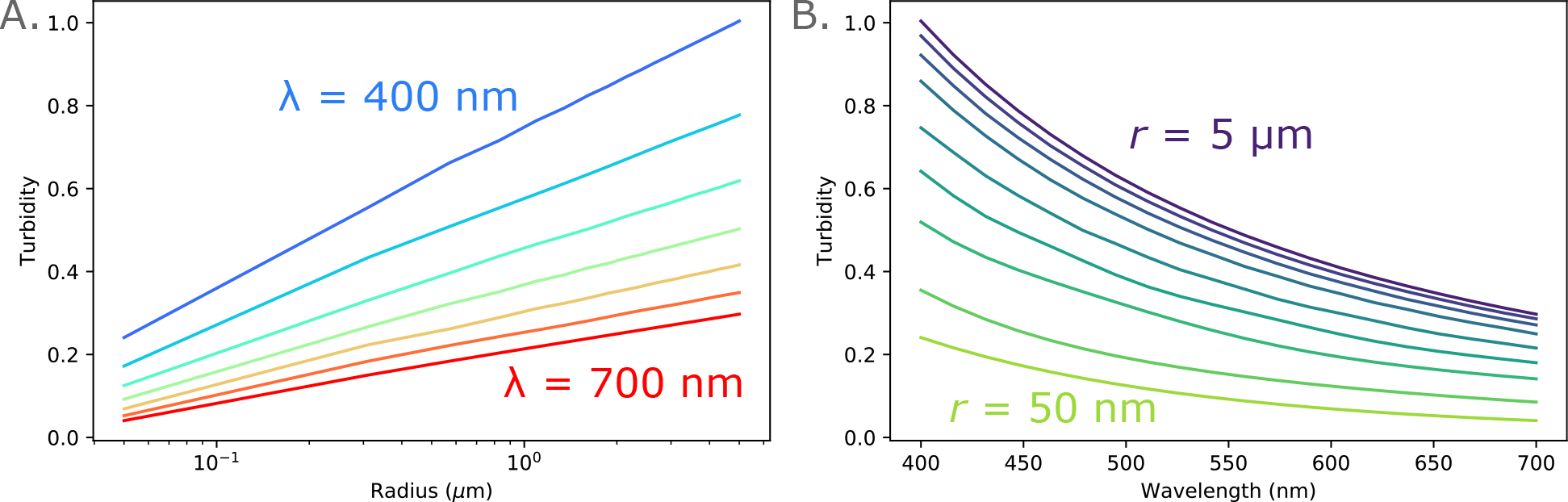
Turbidity of a 5 mM sample of vesicles as a function of A. vesicle radius (λ = 400 nm, 450 nm, 500 nm, 550 nm, 600 nm, 650 nm, 700 nm) and B. wavelength (*r* = 50 nm, 100 nm, 250 nm, 500 nm, 1 *μ*m, 2 *μ*m, 3 *μ*m, 4 *μ*m, 5 *μ*m).

### D. Vesicle lamellarity

Thus far we have only considered scattering from unilamellar vesicles. Experimentally, vesicles often assemble into multilamellar structures so here we determine the effect of lamellarity on sample turbidity. We find that the calculated turbidity of a 5 mM bi-lamellar vesicle suspension is identical to that of a 10 mM unilamellar sample (Fig. 5A). More generally, for a fixed concentration of lipid (5 mM) and wavelength (400 nm), we find that the turbidity *τ_q_* of a sample of *q*-bilayered vesicles is *q*-times more than that of a unilamellar vesicle sample (Fig. 5A) of turbidity *τ*_1_: *τ_q_* = *qτ*_1_. A 5 mM solution of 10-bilayered vesicles will thus scatter like a 50 mM solution of unilamellar vesicles.

**FIG. 5.**
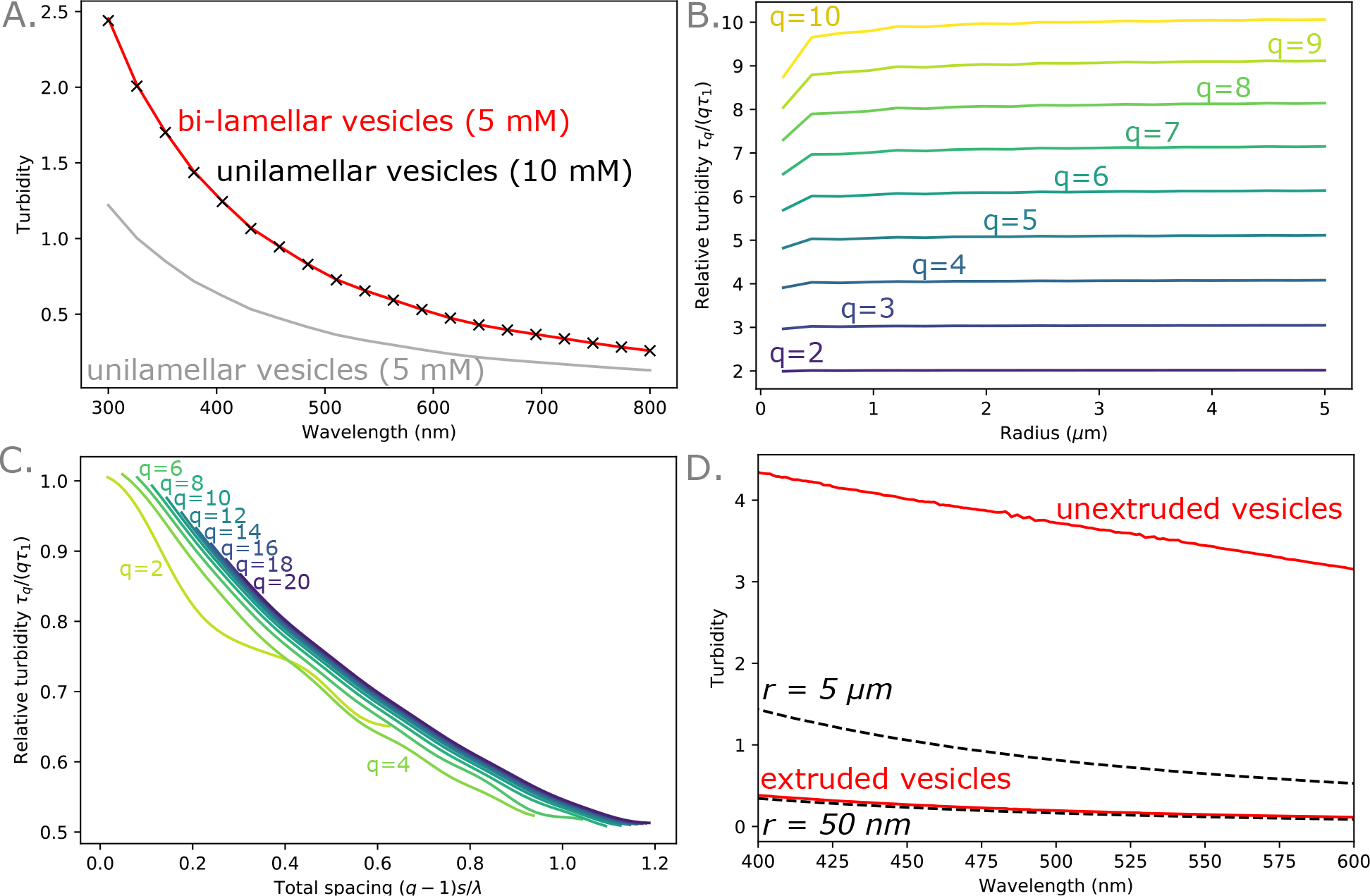
**A**. The calculated turbidities for a 5 mM sample of bi-lamellar vesicles and a 10 mM sample of unilamellar vesicles are identical, and twice that of a 5 mM sample of unilamellar vesicles (*r* = 0.5 *μ*m). **B**. The calculated turbidity of a *q*-bilayered-sample relative to a unilamellar sample is *q*. *λ* = 400 nm. **C**. A sample’s relative turbidity scales with the total spacing between the innermost and outermost membranes (*q* - 1)*s*/λ (*r* = 0.5 *μ*m, λ = 400 nm). **D**. Experimentally measured turbidities of unextruded vesicles and vesicles extruded through 50 nm pores are shown for 5 mM oleic vesicle samples (red solid lines). Calculated turbidities are shown for 5 mM 100-nm-diameter unilamellar vesicle samples and 5 mM 10-*μ*m-diameter unilamellar vesicle samples (black dotted lines).

The strong dependence of the scattering cross section on lamellarity is a consequence of Rayleigh scattering. Because most vesicle membranes are on the order of 5 nm in thickness, the membranes behave as Rayleigh scatterers (at optical wavelengths) for which the scattering cross-section *σ*_sca_ is proportional to *m*^2^, the square of the scatterer’s mass^24^. If two membranes are in close proximity, they will scatter as a single membrane of twice the thickness and hence mass. The scattering cross section of an *q*-bilayered vesicle *σ*_sca,*q*_ is therefore *q*^2^ the scattering cross section of a unilamellar vesicle of the same size *σ*_sca,1_. For a fixed concentration of lipid, the number of vesicles scales inversely with the lamellarity, and so an *q*-bilayered sample is expected to scatter *q*-times more than a unilamellar sample of the same concentration, leading to the same scaling seen as in our calculations (Fig. 5).

We seek to understand how the effect of lamellarity on scattering is weakened with increased intermembrane spacing *s*. For 1-*μ*m-diameter vesicles, we find that while for closely spaced membranes *τ*_*q*_ ~ *qτ*_1_, the ratio *τ_q_/qτ*_1_ decreases from 1 as spacing between the innermost and outermost membranes (*q* − 1)*s* increases (Fig. 5C). τ_1_ is the average turbidity of unilamellar vesicle samples with vesicle radii equalling that of the layers in the multilamellar vesicle. The scaling is similar regardless of the total lamellarity, which suggests that it is the distance between the inner- and outer-most membranes that determines the strength of the dipole coupling.

The strong dependence of turbidity on lamellarity is surprising, and we thus seek to track the turbidity of a sample as it changes from being multilamellar to unilamellar. It is commonly noted that when a milky, heterogeneous vesicle sample is extruded through pores less than 200 nm in radius, the sample will become more transparent^32^. Vesicles are large and multilamellar prior to extrusion, and become small and predominantly unilamellar after extruding through pores smaller than 200 nm in diameter^33^.

Because one of the outcomes of extrusion is to create smaller vesicles, we must attribute part of the decrease in turbidity to the size change. However, our results (Fig. 4) show that for the same concentration of lipid, the turbidity depends only weakly on the vesicle size. We propose that the dominant contribution to the change in vesicle turbidity during extrusion is a change in the lamellarity of the vesicles.

We have measured the turbidities of extruded and unextruded vesicle samples, and also calculated the corresponding turbidities of unilamellar vesicles with our model using the wavelength-dependent refractive index of oleic acid from Jones *et al.*^34^ for *n*_membrane_, and wavelength-dependent refractive index of water from Engen *et al.*^35^. Our results (Fig. 5) show that the extruded samples scatter as expected from the calculated scattering of a sample of 100-nm-diameter unilamellar vesicles. The calculated turbidity for the largest unilamellar vesicles that can occupy the volume (r = 5 *μ*m) is still much smaller than the experimentally measured turbidity for unextruded vesicles, suggesting that the size of vesicles alone can not completely account for the excess scattering. Because extrusion decreases both the size and lamellarity of samples, it is highly likely that multilamellarity is responsible for the excess scattering of unextruded vesicles. For example a tri-lamellar sample of 5 *μ*m vesicles would approximately have the values measured experimentally. However at such high experimentally measured turbidity values, the sample is likely to be highly multiply scattering and our simple model (Eq. 4) is no longer appropriate for direct comparisons.

### E. Presence of aggregates

We also consider cases where an amphiphile does not form membranes in solution, but instead forms aggregates. This can happen when the ionic strength of the solution is too high, there is precipitation (for example divalent cations with fatty acids), the temperature is below the transition temperature of the amphiphile, or in the case of pH-sensitive molecules, the pH is unsuitable. We model aggregates as solid spheres with *n*_contents_ = *n*_lipid_.

We find that if a sample of aggregates is formed, the scattering depends non-monotonically on wavelength and aggregate size (indicative of the aggregates being Mie scatterers^24^) and can dramatically exceed that of vesicles (Fig. 6). The non-monotonic scaling of turbidity with aggregate size means that if the aggregates were to further aggregate, the turbidity of the sample could either increase or decrease depending on the average aggregate size.

**FIG. 6.**
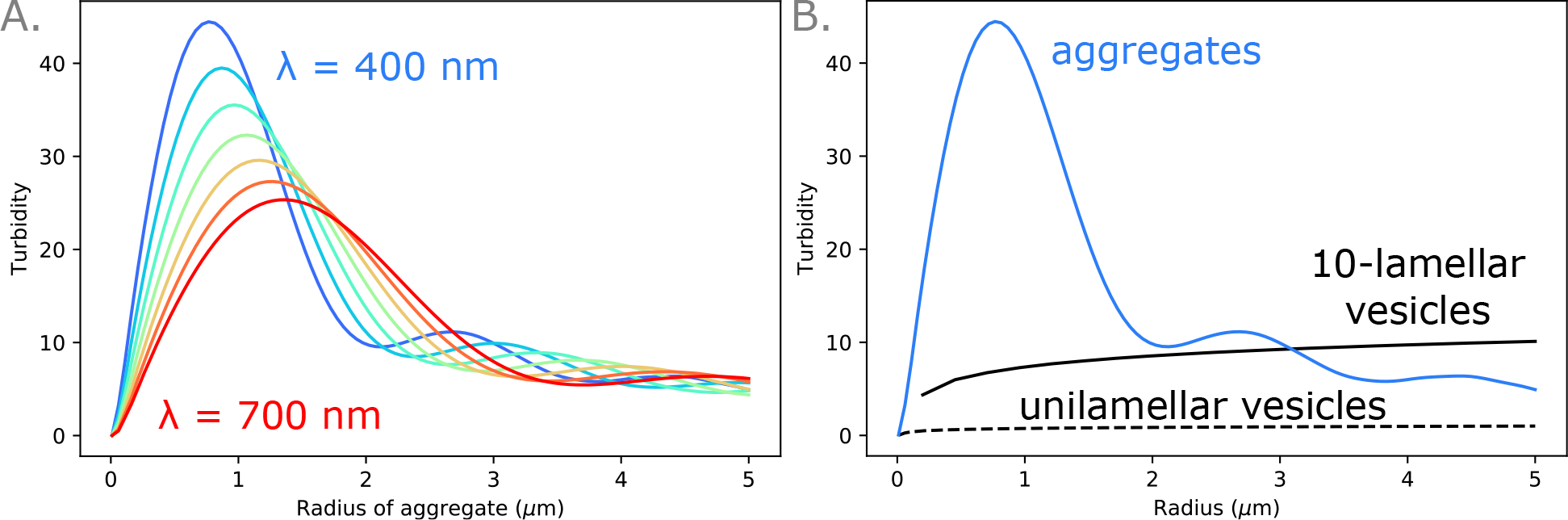
A. Scattering is a non-monotonic function of the aggregate radius, an effect that is typical of Mie scattering. For aggregates smaller than 1 *μ*m, scattering increases with radius. *c* = 5 mM, λ = 400 nm, 450 nm, 500 nm, 550 nm, 600 nm, 650 nm, 700 nm. B. A 5 *μ*m sample of lipid can scatter very differently depending on how the material is arranged. A sample of 1-*μ*m-diameter aggregates scatters more than a sample of 10-bilayered vesicles, which in turn scatters more than a sample of unilamellar vesicles. λ = 400 nm.

Importantly, these results show that even a small contamination of a vesicle suspension with aggregates could dramatically change the turbidity and render such measurements useless. To mitigate the potential confusion, extrusion through small pores, sonication, and adequate mixing can all help reduce the number of aggregates present in a sample.

### F. Effects of scattering towards the detector and multiple scattering

Thus far we have made the assumption that all of the light reaching the detector is unscattered light. However, some vesicles do scatter significantly in the forward direction (0°) and the detector is not a infinitesimal pinhole, but subtends a finite angle (Fig. 7A). Thus here we seek to determine the effect of scattering towards the detector, and when that needs to be taken into account.

**FIG. 7.**
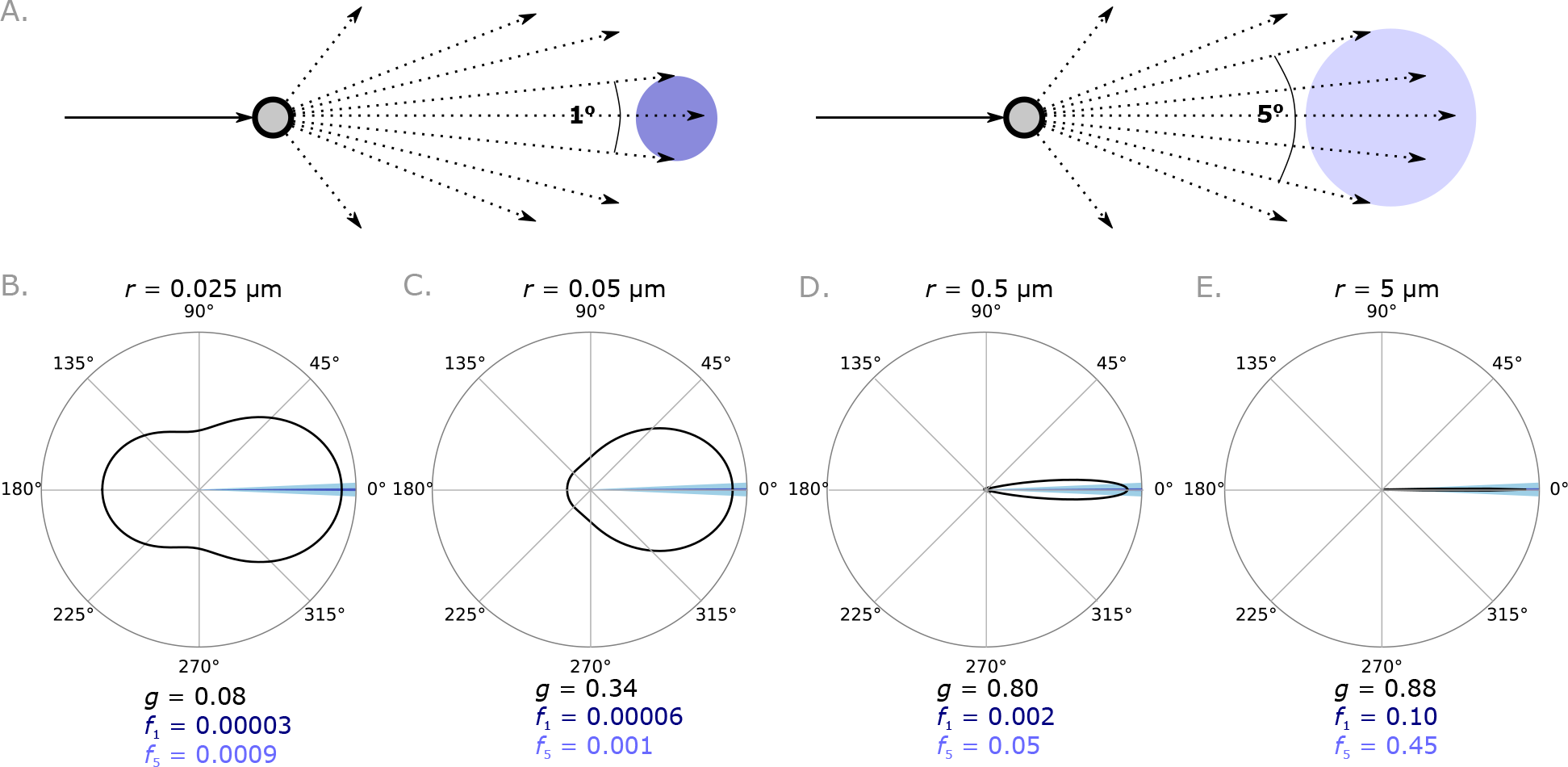
A. Vesicles that scatter forwards can scatter light towards a detector. The fraction of scattered light reaching the detector depends on the detector size. B-D. The intensity of light scattered as a function of angle is shown on polar plots, and the asymmetry parameter *g*, and fraction of scattered light reaching detectors with circular apertures subtending 1° (*f*_1_) and 5° (*f*_5_) are shown. Calculations are done for four different vesicle sizes. Larger vesicles scatter more in the forward direction. λ = 400 nm.

We calculate scattering as a function of angle (scattering phase functions) and in Figure 7B-E show that vesicles larger than 100 nm in diameter scatter significantly in the forward direction. We include the fraction of scattered light that reaches detectors with acceptance angles of 1° (*f*_1_) and 5° (*f*_5_), as well as the asymmetry parameter, *g*, a quantity used to describe the average angle of the scattered light^36^. In general, scatterers with a size much smaller than 1/10 of the wavelength of light will scatter more isotropically; this is indeed true for the smaller vesicles (Fig. 7). Conversely, scatterers that are much larger tend to scatter more light forwards^24^.

To quantify how much extra light is reaching the detector because of forward scattering, we consider two models. The first one models single scattering with the exact phase function and the second one models multiple scattering using an approximation of the phase function^37^.

When considering singly-scattering samples, the intensity of light reaching a detector in the absence of absorption and forward scattering is

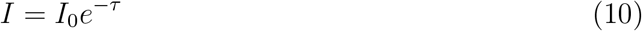

The amount of light that is scattered *I*_*s*_ is hence

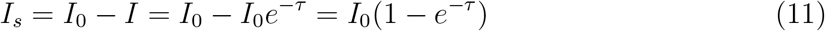

The amount of light that is scattered to a detector that has a circular aperture with acceptance angle *d*° is

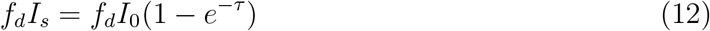

The amount of light *I*_obs_ observed by the detector, including unscattered light, is hence

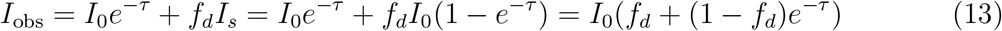

The observed turbidity τ_obs_ is thus

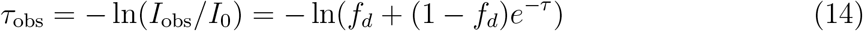

We can therefore use Equation 14 to understand how much the turbidity detected on a spectrophotometer τ_obs_ underestimates the true turbidity *τ* as a result of forward scattering.

We plot τ_obs_/τ in Figure 8 as a function of *f*_*d*_ and τ, with the four vesicle types in Figure 7B-E overlaid. As expected, the experimentally observed τ_obs_ is approximately equal to τ for smaller vesicles because they scatter less strongly in the forward direction, but the disagreement between τ_obs_ and τ grows with increasing *d* and *f*_*d*_. Because τ_obs_/τ depends on the acceptance angle *d* and hence the geometry of the spectrophotometer, it may be difficult to directly compare measurements made on different spectrophotometers. Spectrophotometers are thus suitable for quantifying the turbidity of samples of smaller vesicles; measurements of samples of larger vesicles can severely underestimate τ because of forward scattering.

**FIG. 8.**
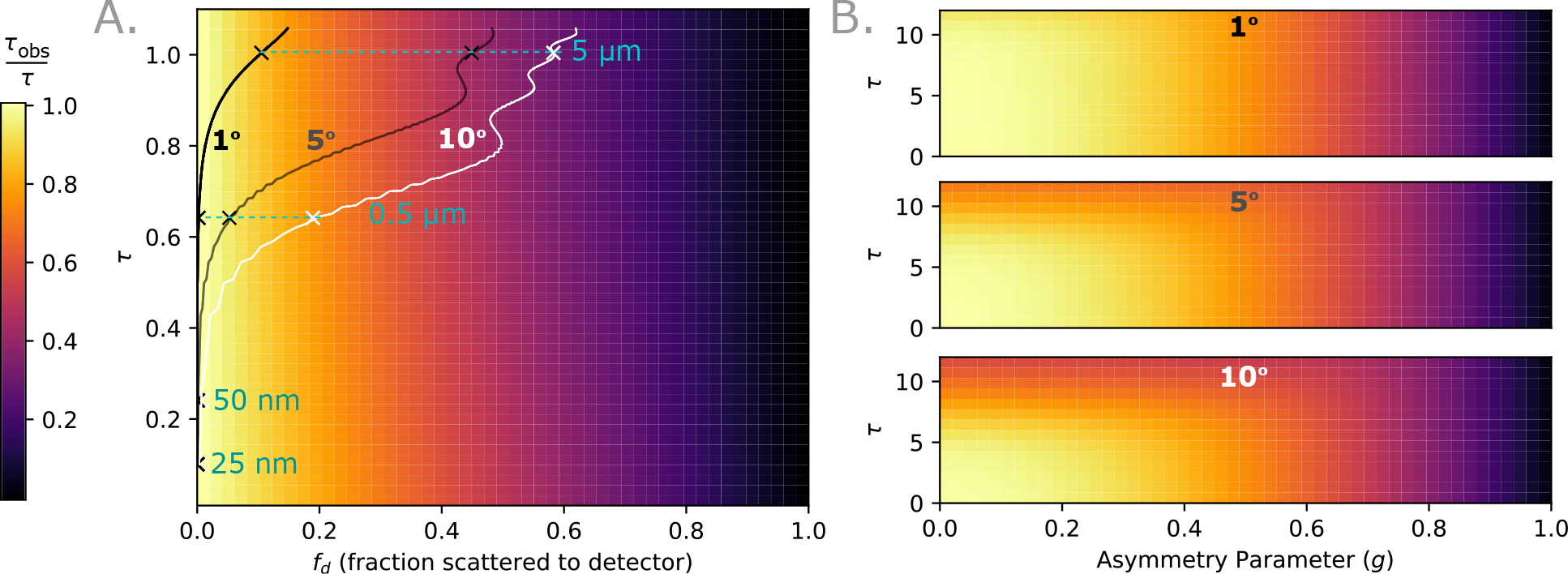
A. τ_obs_/τ is plotted as a function of *f*_*d*_ and τ assuming no scattering in the direction of the detector. The *f*_*d*_ and τ for 1° (black), 5° (grey), and 10° (white) circular apertures are shown for 5 mM samples of vesicles with different radii. A 5 mM sample of the vesicles from Figure 7B-E are marked with blue crosses. λ = 400 nm. B. τ_obs_/τ is plotted as a function of *g* and τ for 1°, 5°, and 10^°^ circular apertures.

We now seek to determine the effect of multiple scattering on the measured turbidity τ_obs_. We use a simple two-stream radiative transfer model^38^ that is more accurate than Equation 4 for large τ, but approximates the phase function^37^. We find that even for small scatterers (*g* ~ 0), τ_obs_/τ can decrease from unity (Fig. 8B) if the sample is concentrated enough. Again, τ_obs_/τ depends on the geometry of the instrument. For further discussions of multiple scattering, we refer the reader to papers that provide accessible overviews of the relevant concept^39–41^.

### G. Implications for experimental design

We have shown how the measured turbidity of a sample depends on the amphiphile concentration *c*, path length *l*, and *p* = (*r*_*i*_, *n*_contents_, *t*, *n*_membrane_}, summarised in Table I. To measure any one parameter with a spectrophotometer, one must control for all of the others. We have also shown the limitations of using spectrophotometers to measure turbidity, with the simplest results to interpret being for a dilute sample of small vesicles.

**TABLE I.** Summary of how τ scales with different parameters.

| Parameter | Explanation | Scaling |
| --- | --- | --- |
| $r$ | vesicle radius | ∼logarithmic |
| $t\Delta n$ | optical path difference | quadratic |
| $\Delta n_{\text{io}}$ or $k$ | $n + ik - n_{\text{medium}}$ | ∼quadratic |
| $k$ | *absorbing contents | ∼quadratic |
| $q$ | lamellarity | linear |
| $c$ | concentration | linear |
| $l$ | path length of light | linear |

We now illustrate how turbidity can be used in conjunction with other tools to gauge information about experimental samples.

#### 1. Experimental results 1. Interpreting turbidity measurements during vesicle formation

During de novo vesicle formation that is triggered by a drop in pH, turbidity can be used to monitor the assembly of vesicles from micelles. In brief, a solution of micelles at high pH is added to a solution buffered at a pH near the pKa of the fatty acids. At this lower pH, vesicles are thermodynamically favoured over micelles. A commonly used assumption is that any increase in scattering of the solutions is because of the fatty acids rearranging into vesicles, and scattering more light. The turbidity of the sample is hence expected to increase over time.

Upon adding a solution of oleic acid micelles to 50 mM bicine (pH 8.1) to a final oleic acid concentration of 5 mM, we find that the turbidity changes non-monotonically with time (Fig. 9). The sample turbidity initially increases to a maximum one day after the sample is first made, then the decreases during the second day. The decrease in turbidity is at first glance surprising.

By monitoring the samples with phase contrast microscopy, we were able to determine that at 1 day, the sample consists mostly of highly-scattering non-spherical aggregates or extremely multilamellar, non-spherical vesicles with little encapsulation volume (Fig. 9). From Figure 6B we can see that for the same concentration of lipid, aggregates and very multilamellar vesicles can scatter a lot more light than oligolamellar vesicles. We therefore attribute the decrease in turbidity to the aggregates disappearing and giant oligolamellar vesicles forming in their place.

**FIG. 9.**
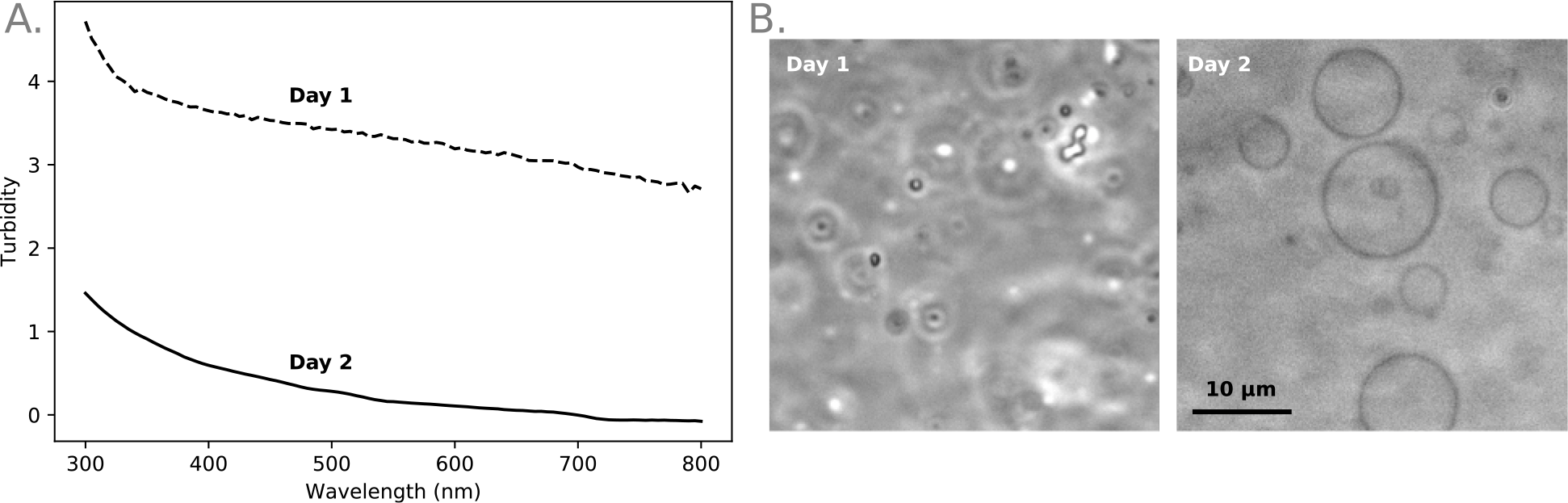
A. A 5 mM oleic acid sample (50 mM bicine, pH 8.1) is highly scattering after one day, but becomes more translucent by day 2. B. Examining the sample with phase contrast microscopy reveals that the sample transitions from being mostly aggregates to mostly oligolamellar.

#### 2. Experimental results 2. Measuring membrane thickness with turbidity

We measure the membrane thickness of oleic acid vesicles by fitting a model containing information about all of the parameters *p* except the thickness *t* to measured sample turbidity. We use the wavelength-dependent refractive index of oleic acid from Jones *et al.*^34^ for *n*_membrane_, and wavelength-dependent refractive index of water from Engen *et al.*^35^. To control for size and lamellarity, we extrude vesicles through 50-nm-diameter pores to achieve an almost completely unilamellar sample^33^, and use the size distribution determined with dynamic light scattering DLS (Malvern Zetasizer Nano C).

Our model fits the data at all wavelengths extremely well. We find that the best-fit thickness for oleic acid is 3.2 nm (s.d. 0.1 nm, n=4). This is in good agreement with cryo-TEM measurements by Namani *et al.*^42^ and simulations by Han^25^. Assuming that at all wavelengths *n*_oleic_ – *n*_palmitoleic_ ¡ 0.01 and *n*_palmitoleic_ – *n*_myristoleic_ ¡ 0.01, the thickness of palmtioleic acid/palmtioleate and myristoleic acid/myristoleate membranes are 2.7-2.9 nm and 2.5-2.7 nm. We show the best-fit results for three vesicle samples in Figure 10. The size distributions used in the model are lognormal, with arithmetic mean and standard deviation taken from DLS measurements (in Fig. 10 2*r* = 124 ± 42 nm for oleic acid, 2*r* = 109 ± 37 nm for palmitoleic acid, and 2*r* = 86 ± 30 nm for myristoleic acid).

**FIG. 10.**
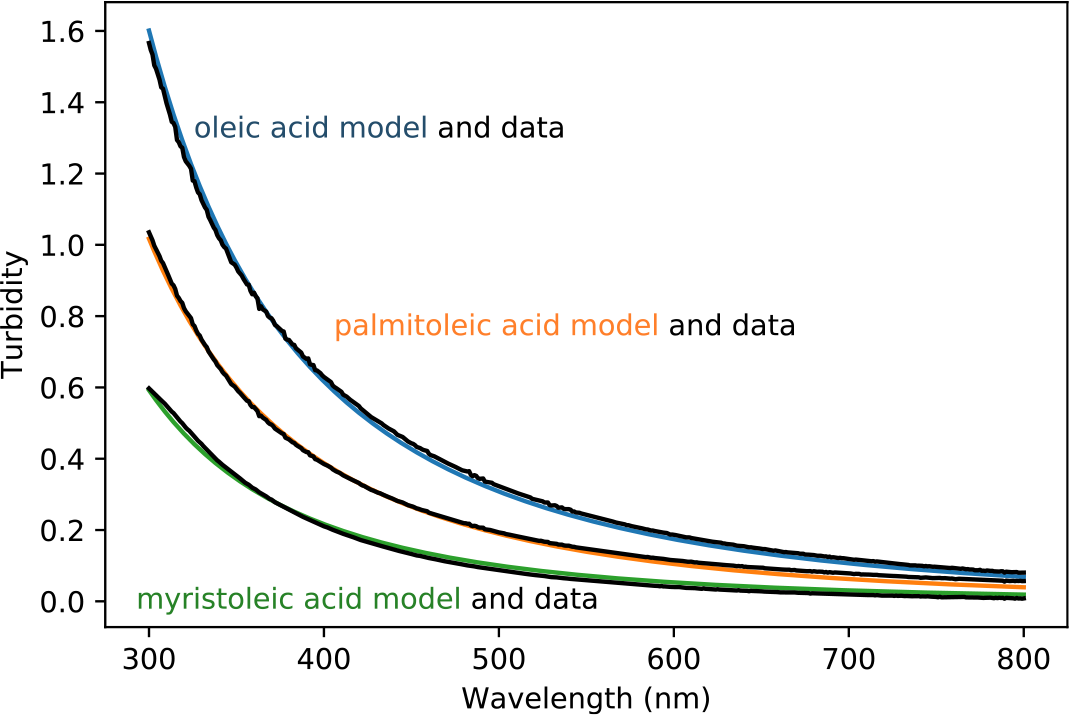
The turbidity of oleic acid, palmitioleic acid, and myristoleic acid vesicles after extrusion are fit well by a model that takes into account the size distributions, concentration, and vesicle composition. The buffer is 200 mM bicine, pH 8.1 for all samples. The concentration of lipids partitioned into the vesicle phase is 8.26 mM minus the critical vesicle concentration from Budin *et al.*^43^.

These results clearly demonstrate that measurements of the parameters *p* can be made if there is knowledge of the rest of *p*.

## IV. DISCUSSION

Using a core-shell model of how vesicles scatter light, we have shown how the turbidity of a vesicle sample changes with lipid refractive index, encapsulated contents, vesicle size, lipid membrane thickness, and lamellarity. Our model is valid when the vesicle solution is in the single scattering regime (turbidity scales linearly with concentration), and there is insignificant scattering in the forward direction. If the vesicle contents have a large refractive index contrast with the medium, or there are many Mie scatterers in solution (e.g. colloidal particles, aggregates), the assumptions that the sample is minimally forward scattering and in the the single scattering regime break down and it becomes difficult to interpret turbidity data in a quantitative manner. As an aid, we provide guidelines for when the observed (τ_obs_) and modelled (τ) turbidity begin to differ significantly. We note that the dependence of τ_obs_ on the detector acceptance angle *d* means that this correction is instrument-dependent. Knowledge of the exact light path within the spectrophotometer is required to make a complete model.

Because the turbidity depends on so many parameters, tools other than a spectrophotometer must be used to understand which parameters are contributing to sample turbidity. For example without microscopy images, it may have been tempting to interpret the high turbidity of the sample in Figure 9 at 1 day as the presence of a high concentration of vesicles. We reveal that in fact the turbidity was because of a high concentration of aggregates, and that seemingly paradoxically, the concentration of vesicles increases only when the turbidity drops.

Finally we showed that with careful experimental design, turbidity can be quite a powerful tool. By using extrusion to constrain lamellarity, and dynamic light scattering to measure the vesicle size distribution, we were able to model how such a vesicle sample would scatter light to measure the membrane thickness of oleic acid vesicles. Our results are in agreement with values in literature, but instead of requiring manual measurements on a cryo-TEM, our method uses equipment easily accessible to those that work routinely with vesicles – an extruder, dynamic light scattering instrument, spectrophotometer, and a computer.

## V. MATERIALS AND METHODS

Micelles are made by dissolving 200 *μ*mol of neat fatty acid oil – oleic acid, palmitoleic acid, or myristoleic acid (NuChek Prep) – in 1.25 equivalents of NaOH (Sigma-Aldrich). The solution is then brought to a volume of 2 mL with Millipore water (18.2 MΩ. cm), vortexed, and left on a test-tube rocker (Speci-Mix) for at least 1 hour to yield a 100 mM micelle solution.

To make the concentrated buffer stock (0.5 M), bicine (Sigma-Aldrich) is dissolved in Millipore water (18.2 MΩ. cm), then titrated to the desired pH with NaOH.

Vesicles are made by mixing the micelle solution, buffer stock, and Millipore water to the desired final concentration in a microcentrifuge tube, vortexing for 5 s, then leaving the sample to agitate overnight on an orbital shaker (GeneMate). All vesicle samples are made with 200 mM bicine except for the giant oligolamellar vesicles, which are made with 50 mM bicine. The extruded samples are extruded with a mini-extruder (Avanti) through a Whatman Nucelopore polycarbonate filter with 50-nm-diameter pores 11 times, and left to tumble on a tube rotator (Labquake) for at least one hour prior to making any measurements.

Turbidity is measured on a spectrophotometer (Cary 60, Agilent). A solution containing just the buffer is used as the blank. The vesicle size distribution is measured with dynamic light scattering on a Zetasizer Nano C (Malvern). Samples are held in UV cuvettes (BRAND).

Refractive indices of solutions at 589 nm were measured on an Abbe refractometer (C10 VEE GEE) at 22°. Glucose, sucrose, and adenosine 5’-monophosphate disodium salt (≥ 99%) were all obtained from Sigma-Aldrich. Concentrations of RNA were determined by spectrophotometry on a Nanodrop 2000C (Thermo Scientific).

Yang’s^22^ core-shell recursive algorithm within HoloPy was used to perform all scattering cross section and phase function calculations. HoloPy is open-source and can be found at https://github.com/manoharan-lab/holopy. For comparison to experimental data, the turbidity τ was calculated for a lognormal distribution of radii using the arithmetic mean and standard deviation measured by DLS.

Radiative transfer calculations were done using the solution to the Eddington approximation taken directly from Shettle and Weinman^38^, incorporating the delta-Eddington approximation from Joseph *et al*.^37^. The aperture is assumed to be completely non-reflecting.

## VI. ACKNOWLEDGMENTS

We thank Dr Lijun Zhou for the RNA oligomers that the refractive index measurements were made with, Dr Derek O’Flaherty for helpful discussions and careful reading of the manuscript, and Dr Daniel Duzdevich, Dr Li Li, Travis Walton, and Dr Victor Lelyveld for feedback on the manuscript. J.W.S. is an Investigator of the Howard Hughes Medical Institute. This work was supported in part by a grant from the Simon Foundation (290363) to J.W.S.. A.W. was supported by the NASA Postdoctoral Program.

